# HISS: Snakemake-based workflows for performing SMRT-RenSeq assembly, AgRenSeq and dRenSeq for the discovery of novel plant disease resistance genes

**DOI:** 10.1101/2022.11.01.514708

**Authors:** Thomas M Adams, Moray Smith, Yuhan Wang, Lynn H Brown, Micha M Bayer, Ingo Hein

## Abstract

**Background:** In the ten years since the initial publication of the RenSeq protocol, the method has proved to be a powerful tool for studying disease resistance in plants and providing target genes for breeding programmes. Since the initial publication of the methodology, it has continued to be developed as new technologies have become available and the increased availability of computing power has made new bioinformatic approaches possible. Most recently, this has included the development of a *k*-mer based association genetics approach, the use of PacBio HiFi data, and graphical genotyping with diagnostic RenSeq. However, there is not yet a unified workflow available and researchers must instead configure approaches from various sources themselves. This makes reproducibility and version control a challenge and limits the ability to perform these analyses to those with bioinformatics expertise.

**Results:** Here we present HISS, consisting of three workflows which take a user from raw RenSeq reads to the identification of candidates for disease resistance genes. These workflows conduct the assembly of enriched HiFi reads from an accession with the resistance phenotype of interest. A panel of accessions both possessing and lacking the resistance are then used in an association genetics approach (AgRenSeq) to identify contigs positively associated with the resistance phenotype. Candidate genes are then identified on these contigs and assessed for their presence or absence in the panel with a graphical genotyping approach that uses dRenSeq. These workflows are implemented via Snakemake, a python-based workflow manager. Software dependencies are either shipped with the release or handled with conda. All code is freely available and is distributed under the GNU GPL-3.0 license.

**Conclusions:** HISS provides a user-friendly, portable, and easily customised approach for identifying novel disease resistance genes in plants. It is easily installed with all dependencies handled internally or shipped with the release and represents a significant improvement in the ease of use of these bioinformatics analyses.

## Background

Single Molecule Real-Time – Association genetics Resistance gene enrichment Sequencing (SMRT-AgRenSeq) is a recently developed approach for the identification of novel disease resistance genes in plants based on association genetics. Briefly, the approach utilises Pacific Biosciences (PacBio) HiFi Resistance gene enrichment Sequencing (RenSeq) reads to assemble reference contigs of a plant (cultivar or wild species) thought to contain the target resistance gene. Following this, a *k*-mer based association genetics approach [1] identifies contigs that are positively associated with the resistance phenotype. This approach was recently used to identify a candidate for the *Pr3* gene controlling resistance to *Puccinia recondita* f. sp. *secalis* in Rye [2]. These analyses can be further extended by adding a final step of graphical genotyping via diagnostic RenSeq (dRenSeq) [3, 4] to assess the presence and absence patterns of identified candidates to reduce the number of genes that require *in planta* testing.

To perform this analysis, a user must currently install numerous pieces of software such as Canu [5], Bowtie2 [6] and Samtools [7]. A user must further clone down software from several GitHub repositories for NLR-Annotator [8] and Association genetics Resistance gene enrichment Sequencing (AgRenSeq) [1]. Finally, users must be familiar with a scripting language such as bash to write scripts and perform the analyses. In this paper, we present HISS (HIgh-throughput SMRT-AgRenSeq-d Snakemake), consisting of three Snakemake workflows constituting: SMRT-RenSeq assembly, AgRenSeq for candidate identification and dRenSeq for candidate validation. These reside in a single GitHub repository which contains files detailing the rules, yaml files for install of software via conda, and redistributes required software not available via conda. This represents a significant increase in ease of use, as a user now simply needs to: install anaconda or miniconda, install snakemake, optionally create a Snakemake run profile using cookiecutter, and download the latest release from our GitHub repository [9]. Snakemake then handles the execution of specific rules and reports to the user any issues with the configuration provided or errors occurring during the run. An example dataset is also provided to allow users to test the workflow on their systems.

## Implementation

HISS contains three Snakemake [10] workflows to perform assembly of HiFi RenSeq reads, SMRT-AgRenSeq [2] to identify novel resistance gene candidates and dRenSeq [3] to validate candidate genes or to determine the presence of known genes (Fig. 1). HISS is distributed with yaml files specifying conda environments to ensure reproducibility and version control of software regardless of the host system. A user simply needs to install either anaconda or miniconda and then install Snakemake into their base environment. All other software dependencies will be installed by the workflows on first run. The workflows themselves consist of a series of rules, which Snakemake uses to produce a directed acyclic graph (DAG) and run the analyses. These workflows perform best on a distributed cluster-style system running a scheduler such as slurm, but they could also be run on a sufficiently well-resourced local machine running a Linux distribution. Users are supplied with template configuration files, which are modified by the user prior to performing the analyses. The separate workflows do not strictly depend on one another. However, we anticipate most users will start with SMRT-RenSeq assembly to produce resistance gene containing contigs for use in the SMRT-AgRenSeq workflow and identify candidate genes for their resistance phenotype of interest. These candidates can then be assessed for presence and absence in a panel of samples via dRenSeq. SMRT-RenSeq assembly requires HiFi reads for accurate reconstruction of nucleotide-binding leucine-rich repeat genes (NLRs). The additional samples used for AgRenSeq and dRenSeq only need short, paired-end Illumina reads, reducing the cost implications of these analyses.

**Fig. 1:**
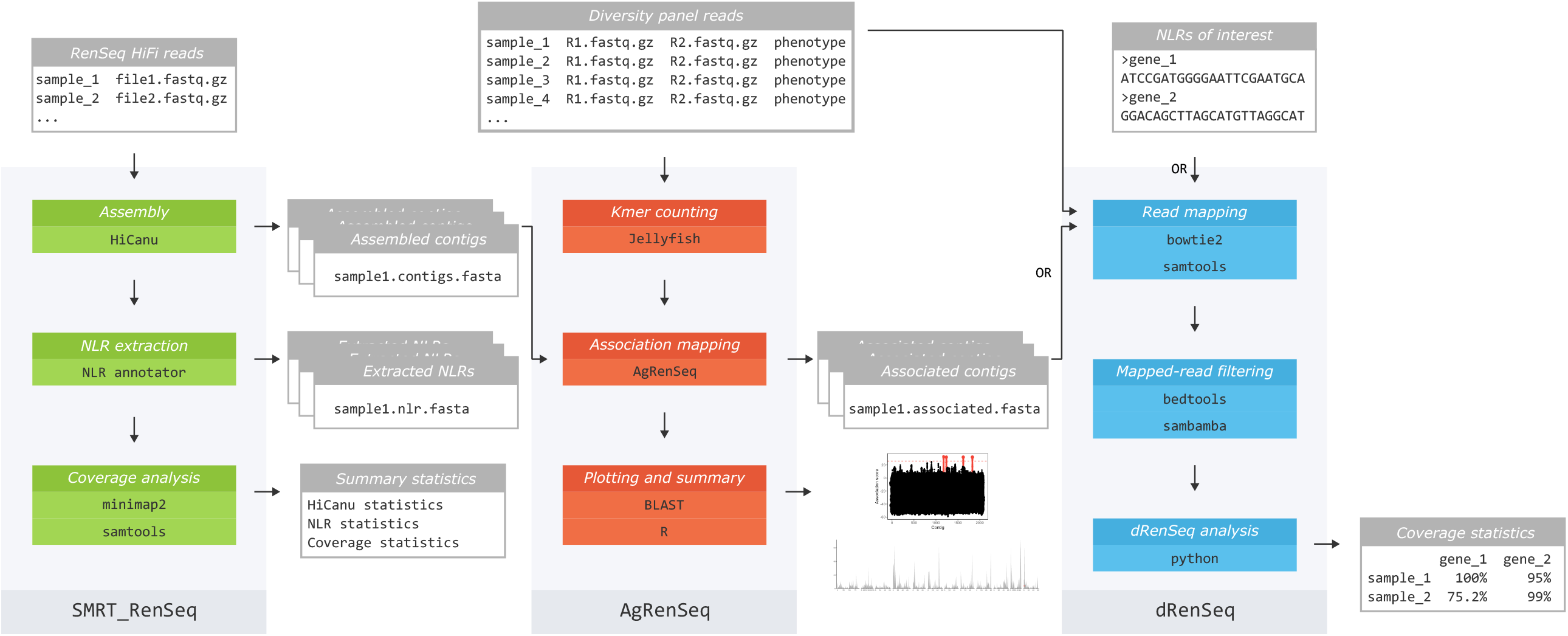
Outline of the Snakemake RenSeq workflows. SMRT-RenSeq (green) assembles RenSeq HiFi reads and produces assembly and NLR summary statistics. AgRenSeq (red) takes a metadata file of diversity panel reads and can use the output of SMRT-RenSeq as a reference for *k*-mer mapping. It outputs highly associated contigs and NLR loci as well as *k*-mer scoring plots and mapping of contigs to a reference genome. dRenSeq (blue) can use a list of NLRs of interest or the output from AgRenSeq to calculate read coverage.

## Workflow summary

All the workflows in HISS begin by parsing a tab-separated values (TSV) file into a pandas dataframe to allow easier wildcarding in subsequent rules.

### SMRT-RenSeq Assembly

#### Read trimming

The first step of this workflow is to perform minimal trimming of sequence from the provided HiFi reads. This step uses cutadapt version 3.5 [11] to remove a provided sequence pattern from the 3’ and 5’ ends of the reads. Users may need to conduct additional trimming depending on what pre-processing has been performed on the raw data prior to starting the workflow.

#### Assembly of reads and basic statistics generation

HiFi reads are assembled into contigs with HiCanu [12] by running Canu version 2.2 [5] with the -pacbio-hifi flag. In addition, useGrid is set to false. When running on a local system, Canu could not utilise grid functionality, whereas on a multi-node cluster system the workflow instead assembles numerous samples in parallel rather than separating a single job across multiple nodes. As targeted enriched reads are used, assembly is also faster than a whole genome sequencing (WGS) assembly, again meaning the benefits of enabling useGrid would be minimal. The user specifies an expected genome size, which is passed to the genomeSize option and represents the expected size of the targeted enrichment space. The maxInputCoverage is also set to 10,000. An early step of Canu is the assessment of the number of bases in the input reads and comparing this to the provided expected assembly size to assess approximate coverage. If this exceeds the maxInputCoverage value, Canu will randomly subsample reads down to the parameters value. Following the completion of assembly, basic assembly statistics are collated by seqfu version 1.9.1 [13] and the number of reads and bases used for assembly are parsed out of the HiCanu assembly report.

#### Identification of putative resistance genes within contigs and assessment of statistics and coverage

NLR-Annotator [8] is used to identify regions in the assembled contigs that contain hallmarks of NLR-like disease resistance genes, based on previously defined MEME motifs [14] using MEME version 5.4.1 [15]. NLR-Annotator version1 and NLR-Parser3 are used with default parameters and custom python code is used to create a summary file of the types of putative NLRs identified across the samples assembled. A custom python script is used to produce a Browser Extensible Data (BED) format file from the NLR-Annotator results. The trimmed HiFi reads are mapped back to the assembled contigs using minimap2 version 2.24 [16] with the map-hifi option. Samtools version 1.13 [7, 17] is used to produce a file detailing the coverage of each putative NLR using the bedcov command. Finally, a custom python script is used to calculate the percentage coverage across the gene and written to a new file. The python scripts are all present in the GitHub repository for the project. A summary of the workflow is provided in Supplemental Figure S1.

### AgRenSeq

#### Read trimming

Illumina RenSeq reads of the diversity panel provided by the user are pre-processed with fastp version 0.23.2 using default options [18]. Reads that have already been trimmed and quality filtered experience minimal data loss.

#### *k-*mer counting and aggregation

For each set of RenSeq reads, *k*-mers are counted via jellyfish version 2.2.10 count [19] with options -C -m 51 -s 1G -t 4. A tab-delimited dump file is created using jellyfish dump with options -L 10 -ct. Dump file paths are aggregated into a single file as a prerequisite for the AgRenSeq *k*-mer presence/absence matrix creation. The created matrix contains presence and absence scores for the identified *k*-mers in each sample.

#### AgRenSeq association scoring

The AgRenSeq matrix is combined with the phenotypic scores specified by the user and the association mapping is performed on each of the RenSeq reference assemblies provided [1]. Each RenSeq assembly is also filtered with NLR-Annotator version 1 [8] with MEME version 5.4.1 [15] as a prerequisite for association mapping, to retain only contigs with NLR genes for further analysis.

#### Result processing and reporting

The result of each AgRenSeq run is provided as a plot of the association scores of each *k*-mer to their respective contig, with the *k*-mers that exceed the predefined association threshold highlighted. To predict the positions within a genome of the contigs that exceed the threshold, RenSeq reference assemblies are mapped to a user-provided genome via BLASTn version 2.13.0 [20] using default options. Highly associated assemblies are highlighted to indicate physical linkage. Plots are created in R version 4.2.2 [21] with the libraries dplyr version 1.0.9 [22] and ggplot2 version 3.3.6 [23]. A summary of the workflow is provided in Supplemental Figure S2.

#### dRenSeq Read trimming

This workflow starts by removing sequencing adaptors provided in user specified fasta files with cutadapt version 3.5 [11]. At this step the reads are also quality-trimmed, retaining a minimum length of 50 bp and using a quality score threshold of 20 at both the 3’ and 5’ ends of reads in both the R1 and R2 file.

#### Align reads to reference FASTA file

The user provided reference FASTA file (e.g., contains NLRs derived from the assembled contigs and that display positive association with the traits) is first indexed with bowtie2 version 2.4.5 [6]. Bowtie2 is again used to align the trimmed reads to the indexed FASTA file, using the sample name as the read group identifier, a user specified minimum mapping score to allow mapping at different mismatch rates, a maximum insert size of 1,000, the --very-sensitive flag, the --no-unal flag to suppress unaligned reads in the output file, the --no-discordant option to keep only concordant alignments and a user-specified maximum number of alignments for each read. The resulting sequence alignment map (SAM) file is then sorted and indexed with samtools version 1.14 [7, 17] to produce a sorted and indexed BAM file. The BAM file produced at the alignment step is filtered with sambamba version 0.8.1 [24] to only keep mappings with the [NM] tag at 0, meaning zero mismatches.

#### Mapping of bait sequences

In order to reduce the chance of false negatives resulting from parts of candidate genes not being present in enriched short reads, the RenSeq bait sequences are used to only assess regions of the genes that would be expected to be sequenced. First, a BLAST database of the reference sequences is created and used to map the bait sequences to the reference FASTA file with BLASTn version 2.13.0 [20]. The user is able to customise the percentage identity and percentage coverage thresholds to match their use case. The biostrings R library version 2.66.0 is used within an R version 4.2.2 script to create a reduced BED file [21, 25]. This BED file is compared to the user provided BED file with bedtools intersect using bedtools version 2.30.0 [26]. A check is run with custom python code to ensure all genes have at least one bait sequence mapped to them.

#### Assess coverage of reference genes with no mismatches

Bedtools [26] is used with the coverageBed command to calculate coverage in the BAM file across all sites in the reference FASTA file. Coverage is calculated across all regions where bait sequences are mapped and written to a single file containing coverage information for all genes in all samples. The file is transposed with pandas to produce a more human-readable coverage file. A summary of the workflow is provided in Supplemental Figure S3.

## Results

HISS is shipped with an example dataset, focusing on the potato *Rx* gene which provides resistance against potato virus X in potato [27] with Gemson as a reference resistant cultivar.

The example workflow began with assembling the HiFi reads of Gemson. This produced an assembly of 3,531 contigs representing 42,004,772 base pairs (bp) with an N50 of 12,421 and an auN of 15,062 [28]. NLR-Annotator [8] predicted 2,527 putative NLR genes spread across 1,799 contigs. This consisted of 1,431 NLRs marked as complete and 578 marked as complete pseudogenes. Including partial NLR predictions brought the total NLR genes lacking the pseudogene tag to 1,724 and an additional 803 with the pseudogenes tag. This assembly was used as a reference for the association genetics workflow in which we assessed 117 potato cultivars: 84 negative for *Rx* resistance and 33 containing the *Rx* resistance gene. With an association threshold of 26, this provided four strong candidate contigs for *Rx*. When analysed by the basic local alignment search tool (BLAST) against version 4.03 of the DM genome [29], the contigs all lay within the same 1Mbp bin on chromosome 12 as expected [27] (Fig. 2). Each of the four contigs had only a single predicted NLR from NLR-Annotator, which were subjected to dRenSeq analysis with the final workflow. Additionally, a panel of previously identified resistance genes from potato, namely: *R1, R2-like, R3a, R3b, R8* and *R9a* [30–35], were added to prevent any samples having zero reads mapping to the reference sequences, as this is flagged as an error by the workflow since it can indicate poor sequencing quality or a lack of mappings. Inspecting the coverage values from the dRenSeq analysis (Supplemental Table S1) showed that except for one sample, all four candidates were 100% represented. Only the sample 15_JHL_140_A1 showed 100% coverage of only one candidate. This candidate resides on tig00001343 and following a BLAST against the national centre for biotechnology information (NCBI) nr/nt database [20, 36] showed a hit with the reference *Rx* sequence at 100% identity and 100% query coverage. The remaining three candidates on tig00001423, tig00002055 and tig00002590 yielded top hits for *Rx*/*Gpa2* which are highly similar in sequence. *Gpa2* encodes an NLR that provides resistance against nematodes, which is physically linked to *Rx* [37]. A BLASTN analysis performed using the alternate sequence of *Gpa2* termed *Nem-Gpa2*^Δ*C292*^ [4] as a query against the three candidates above showed that tig00002055 has a match with 100% identity and 100% query coverage.

**Fig. 2:**
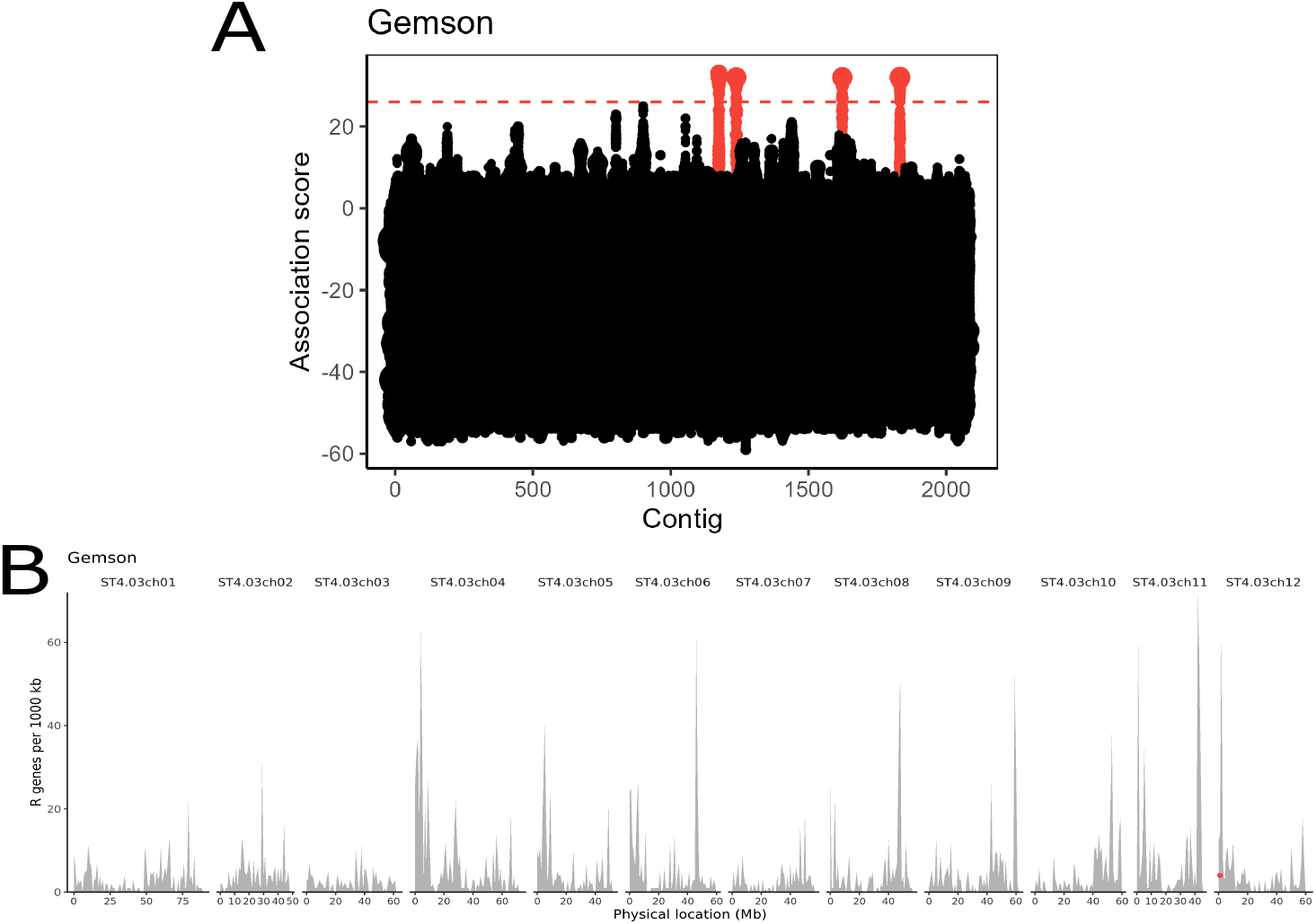
Output plots produced by the Ag-RenSeq snakemake workflow. **A** Dot plot of the analysed contigs in the Ag-RenSeq workflow. Each contig in the assembly is arranged on the x-axis, with *k*-mers plotted on the y-axis. The size of the dot represents the number of *k*-mers. Contigs with *k*-mers above a threshold association score of 26 are highlighted in red. **B** A plot of NLRs by chromosomal location based on the DM reference assembly. The grey bars represent the number of NLRs per 1,000 kb. The red dot represents the location of contigs scored as positive by the association genetics following a BLAST analysis.

## Conclusions

We successfully identified *Rx* and the closely related and physically linked gene *Gpa2* in our SMRT-AgRenSeq analysis. Implementation of dRenSeq in the SMRT-AgRenSeq analysis (coined as SMRT-AgRenSeq-d) identified a rare recombination event in a single sample in our panel that effectively removed three of the four candidates for identification of *Rx*.

Whilst the above example has focused on a known resistance gene, the strength of this combined approach equally applies to identifying elusive genes. Indeed, SMRT-AgRenSeq has recently been used to identify candidate genes for recalcitrant resistance in Rye [2]. With the addition of our dRenSeq workflow, the number of candidates identified by SMRT-AgRenSeq-d can further be refined, reducing both the cost and time required to screen candidate genes *in planta*. As these workflows utilise currently available software and methods, performance improvements will be minimal and limited to the ability of Snakemake to perform multiple rules simultaneously when handling a large sample set, rather than having to wait for all instances of a command to run in a traditional bash script. However, as there are only three points where a user has to start steps of the workflow, this results in less idle time waiting for user input and leads to an improvement in wall clock time elapsed from the start to the end of the analyses. Our modification of dRenSeq to only assess regions covered by bait sequences also reduces the risk of false negatives caused by part of a gene, perhaps separated by a large intron, being absent in the enriched short reads.

## Availability

Project Name: HISS

Project homepage: https://github.com/SwiftSeal/HISS

Operating System(s): Linux

Programming languages: Python, Java, Bash and R

Other requirements: Python 3.10.1, Snakemake 7.12.1, conda 4.13.0, pandas 1.4.3, cookiecutter 2.1.1 (for cluster run profiles only)

License: GNU GPLv3.0

Restrictions to use by non-academics: N/A

Read data for example available at ENA bioprojects: ERP141787 and ERP141790

## List of abbreviations

SMRT-AgRenSeq: Single Molecule Real-Time – Association Genetics Resistance Gene Enrichment Sequencing
SMRT-AgRenSeq-d: Single Molecule Real-Time – Association Genetics Resistance Gene Enrichment Sequencing – diagnostic Resistance gene Enrichment Sequencing
PacBio: Pacific Biosciences
RenSeq: Resistance gene enrichment Sequencing
dRenSeq: diagnostic Resistance gene Enrichment Sequencing
DAG: Directed Acyclic Graph
TSV: Tab-Separated Values
WGS: Whole Genome Sequencing
NLR: Nucleotide-binding, Leucine-rich Repeat
BED: Browser Extensible Data
SAM: Sequence Alignment Map
VCF: Variant Call Format
SNP: Single Nucleotide Polymorphism Bp – Base pairs
BLAST: Basic Local Alignment Search Tool
NCBI: National Centre for Biotechnology Information
HISS: HIgh-throughput Smrt-agrenseq Snakemake

## Declarations

### Authors’ contributions

TMA developed the workflows for SMRT-RenSeq Assembly; MS developed the workflow for AgRenSeq; TMA, YW and LHB developed the workflow for dRenSeq; TMA and MS reviewed the code and contents of the Github repository; MB initially automated the dRenSeq and AgRenSeq approaches; TMA and MS drafted the manuscript; MB and IH conceived the integration of dRenSeq in the workflow and substantively revised the manuscript; IH provided the biological material for the study; all authors reviewed and approved the final manuscript.

## Acknowledgements

We thank Yuk Woon Cheung, Leidy van Rijt and Jannetje C. te Riet for testing the workflows on different datasets and providing helpful feedback and bug reports. We also thank Miles R. Armstrong for the initial development of the dRenSeq protocol. The authors acknowledge the Research/Scientific Computing teams at The James Hutton Institute and NIAB for providing computational resources and technical support for the “UK’s Crop Diversity Bioinformatics HPC” (BBSRC grant BB/S019669/1), use of which has contributed to the results reported within this paper. We also thank John T. Jones for his supervision of Moray Smith’s PhD and for providing comments on the draft.

## Author’s information

TMA and MS contributed equally to this work.

## Figure Legends

Supplemental Table S1: Transposed coverage values table for the example workflow. The reference *R* genes have been removed for clarity. Sample coverage values are colour coded with green representing the highest value and orange representing the lowest value. All samples with the *Rx* gene, except one, show 100% coverage in all four candidates. The 15_JHL_140_A1 represents a rare recombination event in this resistance gene cluster, indicating tig00001343_nlr_1 as a strong candidate for *Rx*.

Supplemental Figure S1: Outline of the SMRT-RenSeq Assembly Snakemake workflow. Input HiFi RenSeq reads are trimmed with cutadapt [11] and assembled in a set of reference contigs with HiCanu [12]. NLR Annotator is used to predict NLR sequences in the contigs [8]. Coverage of the HiFi reads in each NLR prediction is assessed by mapping the HiFi reads back against the assembly with minimap2 [16] and samtools [17].

Supplemental Figure S2: Outline of the AgRenSeq Snakemake workflow. Enriched Illumina reads are trimmed with fastp [18]. The *k*-mers present in these reads are counted with Jellyfish [19]. NLR Annotator is used to identify assembled contigs with signals of NLRs present [8], these contigs are then used to perform the association analysis [1]. The location of candidate contigs in a provided reference genome is assessed by BLAST [20].

Supplemental Figure S3: Outline of the dRenSeq Snakemake workflow. Input enriched Illumina reads are trimmed with cutadapt [11] and aligned to the candidate sequences with bowtie2 [6] and samtools [17]. Bait sequences are mapped to the reference sequences with BLASTn [20] and the regions to be assessed for coverage are selected with the biostrings R package [21, 25]. Bedtools is then used to assess coverage of these regions [26] and a final output file is produced for manual inspection.

